# Reproducibility of the correlative triad among aging, dopamine receptor availability, and cognition

**DOI:** 10.1101/494765

**Authors:** Eric J. Juarez, Jaime J. Castrellon, Mikella A. Green, Jennifer L. Crawford, Kendra L. Seaman, Christopher T. Smith, Linh C. Dang, David Matuskey, Evan D. Morris, Ronald L. Cowan, David H. Zald, Gregory R. Samanez-Larkin

**Affiliations:** Department of Psychology and Neuroscience, Duke University; Center for Cognitive Neuroscience, Duke University; Department of Psychology, Yale University; Center for the Study of Aging and Human Development, Duke University; Department of Psychology, Vanderbilt University; Department of Radiology and Biomedical Imaging, Yale University; Department of Psychiatry, Yale University; Department of Biomedical Engineering, Yale University; Department of Psychiatry and Behavioral Sciences, Vanderbilt University School of Medicine; Department of Radiology and Radiological Sciences, Vanderbilt University Medical Center

**Keywords:** cognition, working memory, aging, dopamine, PET

## Abstract

The evidence that dopamine function mediates the association between aging and cognition is one of the most cited findings in the cognitive neuroscience of aging. However, few and relatively small studies have directly examined these associations. Here we examined correlations among adult age, dopamine D2-like receptor (D2R) availability, and cognition in two cross-sectional studies of healthy human adults. Participants completed a short cognitive test battery and, on a separate day, a PET scan with either the high-affinity D2R tracer [^18^F]Fallypride (Study 1) or [^11^C]FLB457 (Study 2). Digit span, a measure of short-term memory maintenance and working memory, was the only cognitive test for which dopamine D2R availability partially mediated the age effect on cognition. In Study 1, age was negatively correlated with digit span. Striatal D2R availability was positively correlated with digit span controlling for age. The age effect on digit span was smaller when controlling for striatal D2R availability. Although other cognitive measures used here have individually been associated with age and D2R availability in prior studies, we found no consistent evidence for significant associations between low D2R availability and low cognitive performance on these measures. These results at best only partially supported the correlative triad of age, dopamine D2R availability, and cognition. While a wealth of other research in human and non-human animals demonstrates that dopamine makes critical contributions to cognition, the present studies suggest caution in interpreting PET findings as evidence that dopamine D2R loss is a primary cause of broad age-related declines in fluid cognition.

## Introduction

There is considerable evidence from hundreds of studies over the past half century that fluid cognitive abilities decline over the course of the adult life span (Horn & Cattell, 1967; Park & Schwarz, 2000). These fluid cognitive abilities include processing speed, working memory, executive control, and episodic memory. Over the past few decades, functional and structural brain imaging have been used to identify the aspects of neural decline that account for these behavioral losses (Cabeza, Nyberg, & Park, 2016; Grady, 2012; Jagust, & D’Esposito, 2009).

Perhaps the earliest and most consistently reported adult age differences identified in human neuroimaging studies were reductions in aspects of dopamine function. Nearly 100 studies using PET or SPECT imaging have been conducted over the past thirty years documenting lower dopamine receptor and transporter availability in older compared to younger adults. According to a recent meta-analysis (Karrer et al., 2017), adult age is strongly and negatively correlated with D1-like receptor availability (r = –.77), D2-like receptor (D2R) availability (r = –.56), and dopamine transporter availability (r = –.68). All of these measures of dopamine targets (receptors, transporters, or relevant enzymes) have been associated with fluid cognitive abilities (e.g., episodic memory, working memory, cognitive control, psychomotor speed, motor speed) in aging adults (Bäckman, Lindenberger, Li, & Nyberg, 2010).

By the late 1990s and early 2000s, there was evidence for a “correlative triad” suggesting that individual differences in dopaminergic status (assessed with PET/SPECT imaging of dopamine targets), but not chronological age, explain age-related deficits in fluid cognition (Volkow et al., 1998, Bäckman et al., 2000). The implication was that dopaminergic decline makes a critical contribution to fluid cognitive decline with age. The theory is one of the most highly cited in the neuroscience and psychology of aging. However, there are a number of limitations of the existing empirical studies. For instance, although some of the foundational studies included participants across much of the adult life span (Bäckman et al., 2000: ages 21-68, Volkow et al., 1998: ages 24-86) and used the same tracer, [^11^C]Raclopride, each study relied on measures tapping different cognitive functions to provide evidence for the correlative triad. Bäckman and colleagues (2000) reported associations between D2R and perceptual speed and episodic memory whereas Volkow and colleagues (1998) reported associations between D2R and motor and executive function. As can be seen in Table 1, mixed findings, inconsistent reporting of results, and small sample sizes across these studies make evaluation of the reproducibility of these effects difficult. Potential meta-analysis of the previous empirical work is complicated by the inconsistent use of cognitive tests and/or selective reporting of only statistically significant dopamine-cognition associations (Karrer, 2017).

**Table 1.**
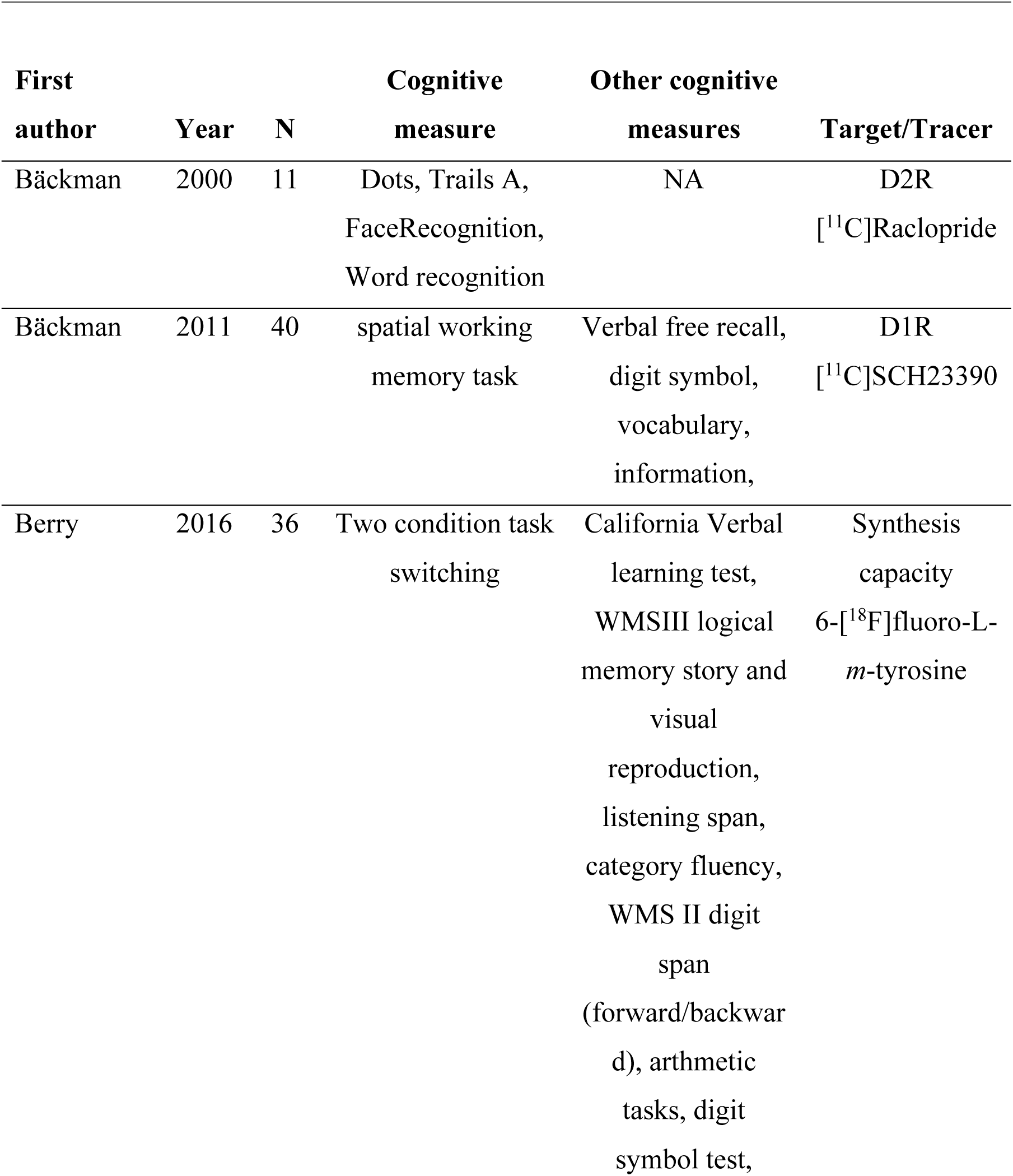
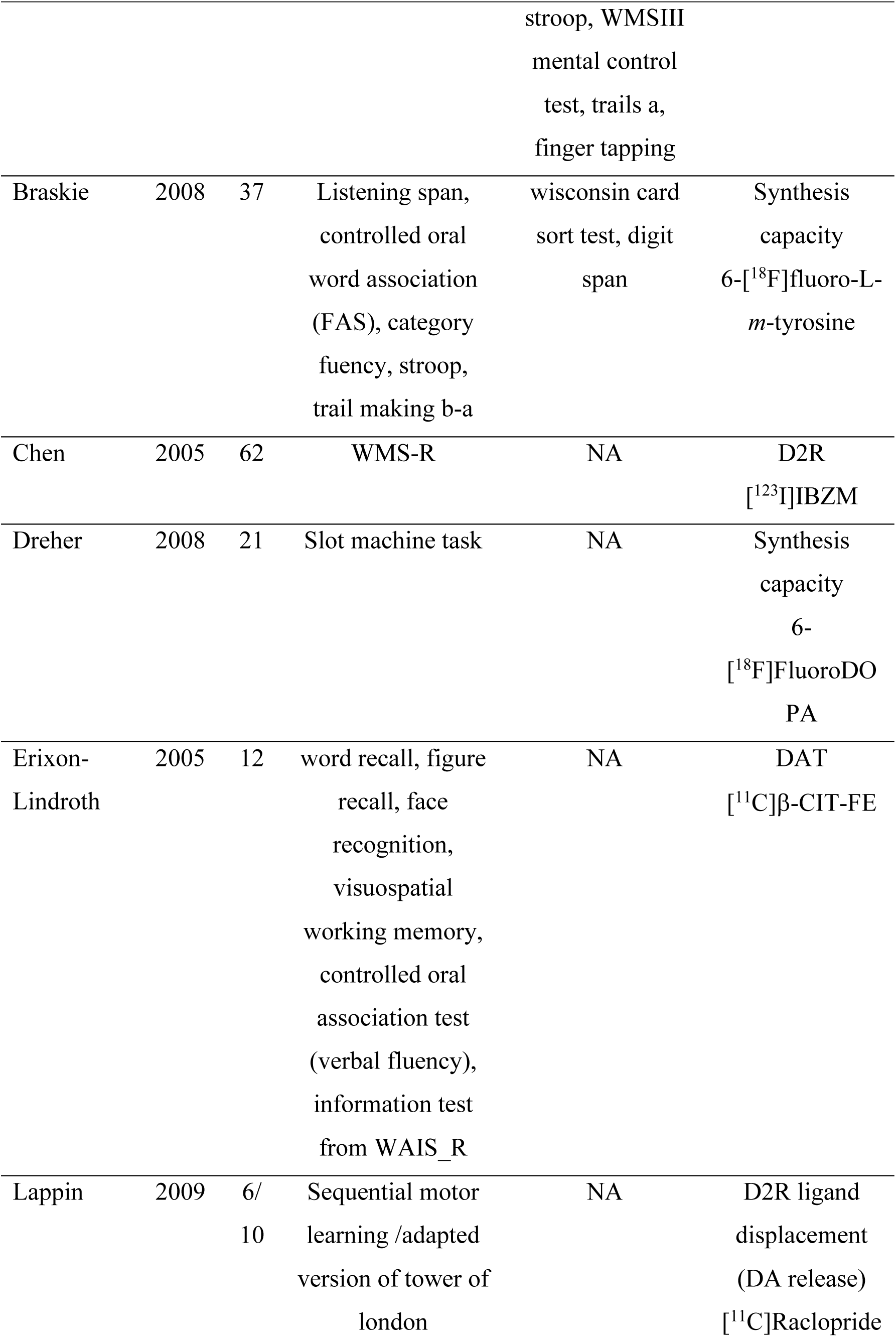
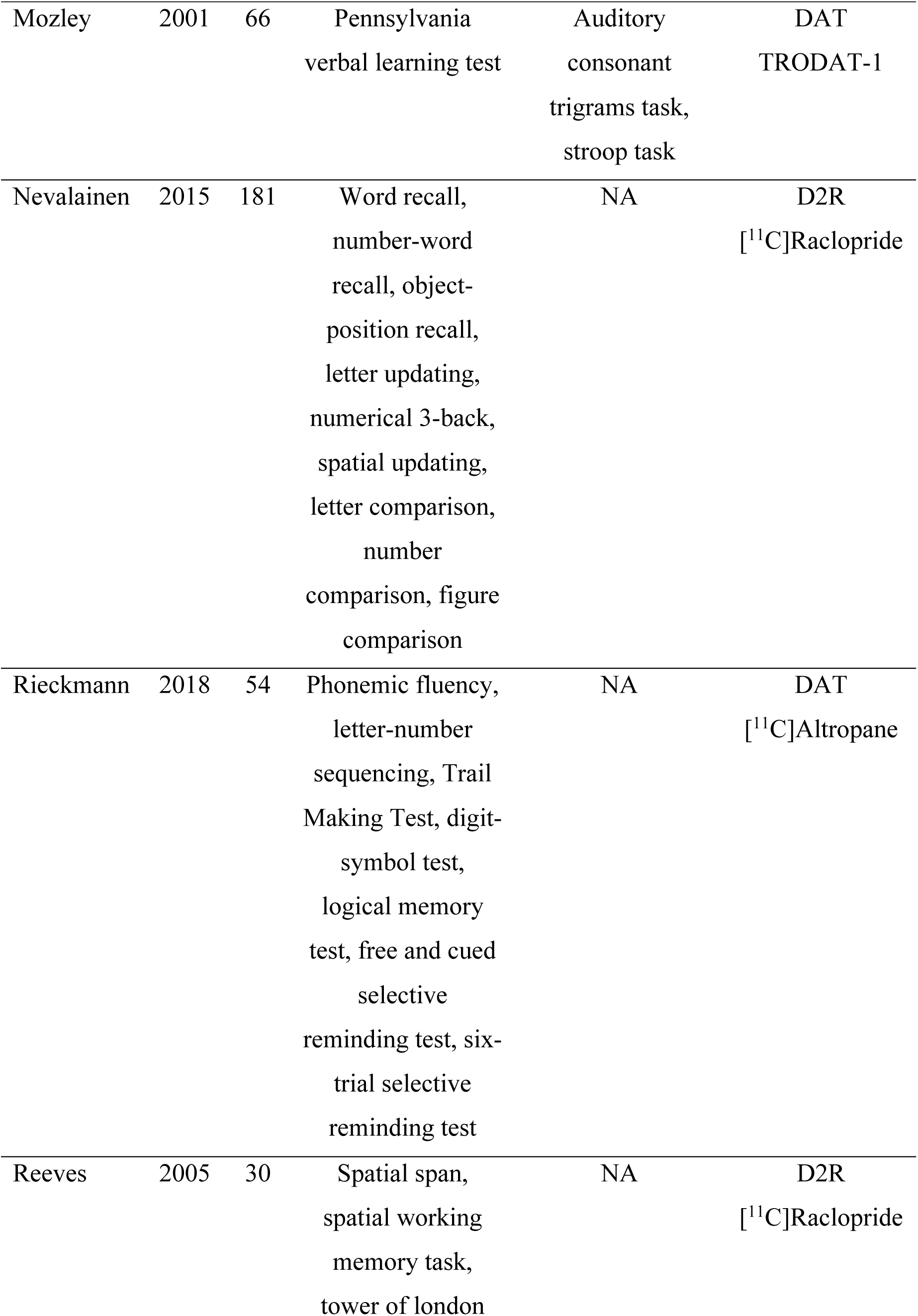
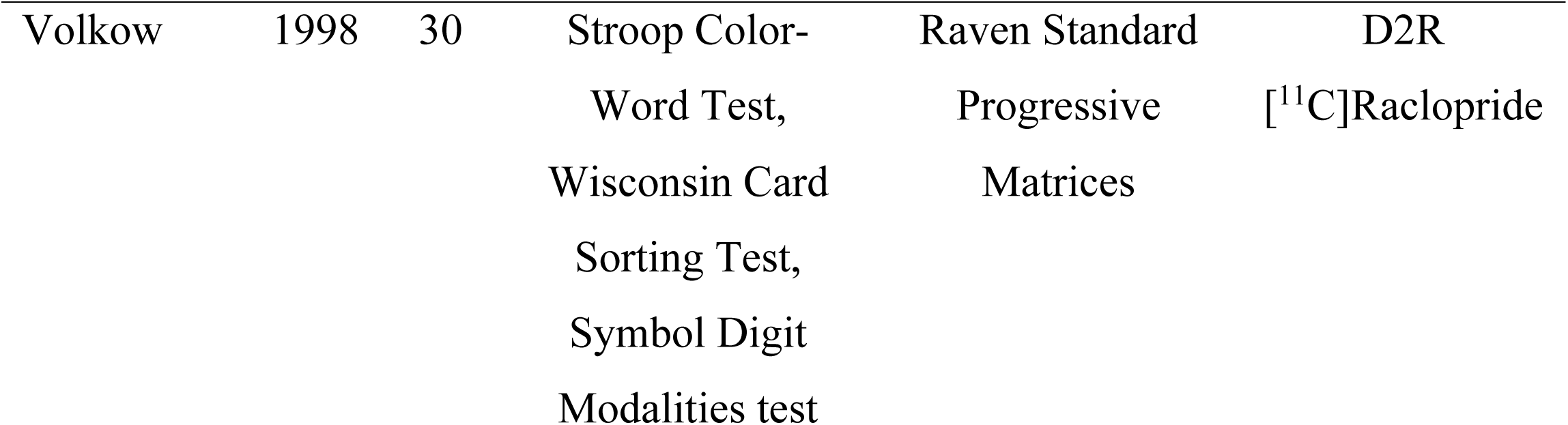
Summary of previous studies investigating aging, dopamine targets (receptors, transporters, or relevant enzymes), and cognition. D1R = dopamine D1-like receptors; D2R = dopamine D2-like receptors; DAT = dopamine transporter.

| First author | Year | N | Cognitive measure | Other cognitive measures | Target/Tracer |
| --- | --- | --- | --- | --- | --- |
| Bäckman | 2000 | 11 | Dots, Trails A, FaceRecognition, Word recognition | NA | D2R<br>[<sup>11</sup>C]Raclopride |
| Bäckman | 2011 | 40 | spatial working memory task | Verbal free recall, digit symbol, vocabulary, information, | D1R<br>[<sup>11</sup>C]SCH23390 |
| Berry | 2016 | 36 | Two condition task switching | California Verbal learning test, WMSIII logical memory story and visual reproduction, listening span, category fluency, WMS II digit span (forward/backward), arithmetic tasks, digit symbol test, stroop, WMSIII<br>mental control<br>test, trails a,<br>finger tapping | Synthesis capacity<br>6-[<sup>18</sup>F]fluoro-L-*m*-tyrosine |
| Braskie | 2008 | 37 | Listening span,<br>controlled oral<br>word association<br>(FAS), category<br>fluency, stroop,<br>trail making b-a | wisconsin card<br>sort test, digit<br>span | Synthesis<br>capacity<br>6-[<sup>18</sup>F]fluoro-L-<br>*m*-tyrosine |
| Chen | 2005 | 62 | WMS-R | NA | D2R<br>[<sup>123</sup>I]IBZM |
| Dreher | 2008 | 21 | Slot machine task | NA | Synthesis<br>capacity<br>6-<br>[<sup>18</sup>F]FluoroDO<br>PA |
| Erixon-Lindroth | 2005 | 12 | word recall, figure<br>recall, face<br>recognition,<br>visuospatial<br>working memory,<br>controlled oral<br>association test<br>(verbal fluency),<br>information test<br>from WAIS_R | NA | DAT<br>[<sup>11</sup>C]β-CIT-FE |
| Lappin | 2009 | 6/<br>10 | Sequential motor<br>learning /adapted<br>version of tower of<br>london | NA | D2R ligand<br>displacement<br>(DA release)<br>[<sup>11</sup>C]Raclopride |
| Mozley | 2001 | 66 | Pennsylvania<br>verbal learning test | Auditory<br>consonant<br>trigrams task,<br>stroop task | DAT<br>TRODAT-1 |
| Nevalainen | 2015 | 181 | Word recall,<br>number-word<br>recall, object-<br>position recall,<br>letter updating,<br>numerical 3-back,<br>spatial updating,<br>letter comparison,<br>number<br>comparison, figure<br>comparison | NA | D2R<br>[<sup>11</sup>C]Raclopride |
| Rieckmann | 2018 | 54 | Phonemic fluency,<br>letter-number<br>sequencing, Trail<br>Making Test, digit-<br>symbol test,<br>logical memory<br>test, free and cued<br>selective<br>reminding test, six-<br>trial selective<br>reminding test | NA | DAT<br>[<sup>11</sup>C]Altropane |
| Reeves | 2005 | 30 | Spatial span,<br>spatial working<br>memory task,<br>tower of london | NA | D2R<br>[<sup>11</sup>C]Raclopride |
| Volkow | 1998 | 30 | Stroop Color-Word Test,<br>Wisconsin Card<br>Sorting Test,<br>Symbol Digit<br>Modalities test | Raven Standard<br>Progressive<br>Matrices | D2R<br>[<sup>11</sup>C]Raclopride |

Only 10 out of 95 previously published datasets on adult age effects on dopamine targets (receptors, transporters, or relevant enzymes) using PET or SPECT reported cognitive associations (Karrer et al., 2017). Ten papers would be sufficient for meta-analysis, but almost none of the papers report associations with the same cognitive measures. The reported associations between different measures of dopamine function and cognitive function spanned several domains such as cognitive control (Lappin et al., 2009), episodic memory (Bäckman et al., 2000), or working memory (Bäckman et al., 2011), but the same cognitive measures were not reported across studies. Recent studies have questioned and begun to revise the broad claims of the original correlative triad hypothesis (Nyberg et al., 2016, Lövdén et al., 2017, Rieckmann et al., 2018, Karalija et al, 2019, Salami, et al., 2019). For example, third variables such as education, lifestyle, or genetics have been implicated as potential critical moderators of dopamine-cognition relationships with aging (Lövdén et al., 2017, Karalija et al., 2019). Thus, given the assumptions that dopaminergic loss underlies fluid cognitive deficits, or may even be a biomarker for cognitive decline, it is important to establish the replicability of the effects.

Here, in two adult life-span studies at different sites using different tracers, with complementary coverage of cortical and subcortical regions, we investigated whether adult age differences in standard neuropsychological measures of fluid cognition were due to individual differences in D2R availability.

## Method

Two studies at two different universities examined the replicability of associations between age, D2R availability, and cognitive test performance.

### Participants

#### Study 1

Healthy participants were recruited from the greater Nashville community for PET scans performed at Vanderbilt University. Study 1 consisted of 83 adults (ages 22 to 83 years, mean=49.80, SD=17.68, 35 males). See Supplementary Figure 1 for age histogram. Participants were medically and mentally healthy and were assessed by a physical examination, comprehensive metabolic panel, complete blood count, EKG, and interviews of medical and psychiatric history. Inclusion criteria were the following: no illicit drug use in last 2 months, no use of any psychotropic medication in last 6 months, no current uncontrolled medical condition such as neurological, cardiovascular, endocrine, renal, liver, or thyroid pathology, no history of neurological or psychiatric disorders, no current tobacco or nicotine use, alcohol consumption no greater than 8 ounces of whiskey (∼5 standard alcoholic drinks)/week, and no claustrophobia or other MRI contraindications. Females had negative pregnancy tests at intake and on the day of the scan. Approval for the Study 1 protocol was obtained from the Vanderbilt University Human Research Protection Program and the Radioactive Drug Research Committee and all participants completed written informed consent.

#### Study 2

Healthy participants were recruited from the greater New Haven community for PET scans performed at Yale University. Study 2 consisted of 37 adults (ages 26 to 79 years, mean=47.81, SD=16.93, 17 males). See Supplementary Figure 2 for age histogram. Participants were medically and mentally healthy and were assessed by a physical examination, comprehensive metabolic panel, complete blood count, EKG, and interviews of medical and psychiatric histories. Inclusion criteria were the following: no prescription or illicit drug use, no history of tobacco or nicotine use, no current uncontrolled medical conditions such as cardiovascular, endocrine, renal, liver, or thyroid pathologies, no history of neurological or psychiatric disorders, alcohol consumption no greater than 8 ounces of whiskey (∼5 standard alcoholic drinks)/week, and no claustrophobia or other MRI contraindications. Females had negative pregnancy tests at intake and on the day of the scan. Approval for the Study 2 protocol was obtained from the Yale University Human Investigation Committee and the Yale University Hospital Radiation Safety Committee and all participants completed written informed consent.

### Cognitive measures

Four cognitive tests were collected in both studies. A relatively brief neuropsychological battery was selected to characterize age differences in common measures of aspects of fluid cognition that had been previously correlated with D2R. All measures were included based on literature indicating that these functions decline with age (Hicks & Birren, 1970; Park & Schwarz, 2000; Salthouse, 2004; Buckner, 2004) and have been linked to prefrontal function or DA modulation (West, 1996; Luciana, Depue, Arbisi, & Leon, 1992; Rubin, 1999). The Trail-Making Test (TMT) measures speed of processing, visual search, and cognitive flexibility (Crowe, 1998). The test includes two parts. In Part A, participants draw lines connecting numbers in numerical order (1-2-3-4) while an experimenter records time to completion.

Completion time on Part A is used as a measure of psychomotor speed and visual search. Part B includes similar psychomotor and search demands but also includes alternation between numbers and letters (e.g., 1-A-2-B-3-C). The difference between these scores (B – A) is used as a measure of cognitive flexibility. Three measures of memory from the Wechsler Memory Scale WMS-III, two that assessed short-term maintenance (Digit Span) and working memory (Digit Span, Letter-Number Sequencing) and one that assessed longer-term memory (Verbal Paired Associates Delayed Recall) were also collected in both studies. The total score of forward plus backward digit span was used for analyses. In the forward digit span task, participants repeat a list of numbers as they were presented by the experimenter. In the backward digit span task, participants repeat a list of numbers in the reverse order. In the letter-number sequencing task, participants must reorder increasingly longer strings of letters and numbers in alphanumeric order. In the verbal paired associates task, participants learn a series of two-word associations which they recall four times immediately after presentation by the experimenter. In the delayed recall portion of the task, participants repeat the paired associates after an extended delay. Only the delayed recall portion of the verbal paired associates task was used for analysis.

### PET Acquisition

#### Study 1

[^18^F]Fallypride, (S)-N-[(1-allyl-2-pyrrolidinyl)methyl]-5-(3[^18^F]fluoropropyl)-2,3-dimethoxybenzamide (hereafter referred to as Fallypride), was produced in the radiochemistry laboratory attached to the PET unit at Vanderbilt University Medical Center, following synthesis and quality control procedures described in US Food and Drug Administration IND 47,245. Prior to the PET scan, T1-weighted magnetic resonance (MR) images (TFE SENSE protocol; Act. TR=8.9 ms, TE=4.6 ms, 192 TFE shots, TFE duration=1201.9 s, FOV=256×256 mm, voxel size=1×1×1 mm) were acquired on a 3T Philips Intera Achieva whole-body scanner (Philips Healthcare, Best, The Netherlands). PET data were collected on a GE Discovery STE (DSTE) PET scanner (General Electric Healthcare, Chicago, IL, USA). Serial scan acquisition was started simultaneously with a 5.0 mCi (185 MBq; average = 5.06 mCi, SD = 0.23) slow bolus injection of the dopamine D2/3 tracer Fallypride (median specific activity: 9.24 mCi/nmol). CT scans were collected for attenuation correction prior to each of the three emission scans, which together lasted approximately 3.5 hours with two breaks for participant comfort. Acquisition times for the dynamic PET scans have been reported previously (Smith et al., 2016). After decay correction and attenuation correction, PET scan frames were corrected for motion using SPM8 (Friston et al., 1995) with the 20^th^ dynamic image frame of the first series serving as the reference image. The realigned PET frames were then merged and re-associated with their acquisition timing information in PMOD (PMOD Technologies, Zurich, Switzerland) ‘s PVIEW module to create a single 4D file for use in PMOD’s PNEURO tool for further analysis (see below).

#### Study 2

[^11^C]FLB 457, 5-bromo-N-[[(2S)-1-ethyl-2-pyrrolidinyl]methyl]-3-methoxy-2-(methoxy-^11^C) benzamide (hereafter referred to as FLB), was synthesized as previously described by Sandiego et al. (2015). PET scans were acquired on the High Resolution Research Tomograph (HRRT; Siemens Medical Solutions, Knoxville, TN, USA). FLB (median specific activity: 7.80 mCi/nmol) was injected intravenously as a bolus (315 MBq; average = 8.62 mCi, SD = 2.03) over one minute by an automated infusion pump (Harvard Apparatus, Holliston, MA, USA). Prior to each scan a six-minute transmission scan was performed for attenuation correction. Dynamic scan data were acquired in list mode for 90 min following the administration of FLB and reconstructed into 27 frames (6 × 0.5 mins, 3 × 1 min, 2 × 2 mins, 16 × 5 mins) with corrections for attenuation, normalization, scatter, randoms, and dead time using the MOLAR (Motion-compensation OSEM List-mode Algorithm for Resolution-Recovery Reconstruction) algorithm (Carson, Barker, Liow, & Johnson, 2003). Event-by-event motion correction (Jin et al., 2013) was applied using a Polaris Vicra optical tracking system (NDI Systems, Waterloo, Canada) that detects motion using reflectors mounted on a cap worn by the participant throughout the duration of the scan. Prior to the PET scan, T1-weighted magnetic resonance (MR) images (MPRAGE protocol; TR=2.4 s, TE=1.9 ms, FOV=256×256 mm, voxel size=1×1×1 mm) were acquired on a 3T Trio whole-body scanner (Siemens Medical Systems, Erlangen, Germany).

### PET data processing

Binding potential was estimated relative to non-displaceable tracer (BP_ND_) within ROIs using PMOD’s PNEURO module. The Hammers atlas (Hammers et al., 2003) available internally in PNEURO and PNEURO’s deep nuclei option was used to parcellate each participant’s grey matter as determined from a segmentation of their T1 MRI image into 30 bilateral cortical areas (including amygdala and hippocampus), 5 bilateral deep nuclei (caudate, putamen, ventral striatum, thalamus, and pallidum), plus bilateral cerebellum, and brainstem. After parcellation, the MRI and PET data were co-registered via a rigid matching procedure based on the normalized mutual information criterion and PET data was resampled to the MRI space (1×1×1mm). Accurate co-registration is critical for the partial-volume correction methods (described later) (Hutton et al., 2013). To check registration quality, we calculated quality control metrics using the PFUS module in PMOD 4.0 for a subset of 42 participants including the oldest 10 participants in each study, since older participants tend to be the hardest to co-register due to grey matter loss with age. The average Dice coefficient between PET data warped to MRI space and each participant’s own anatomical MRI data was 0.83±0.05. The Dice coefficient measures the ratio between the number of true positives (overlapping voxels) compared to the number of true positives (overlapping voxels) plus the number of false positives (non-overlapping voxels). This suggests that registration methods were successful and consistent across the sample. We also tested for age differences in the other registration quality control metrics provided by PFUS (sensitivity, specificity, Jaccard index) across the subsamples and found no relationships between age and any of these registration quality metrics.

The PNEURO parcellation ROIs were then used to extract time activity curves (TACs) from the PET data before modeling. These TACs were then fitted using a two-compartment simplified reference tissue model (Lammertsma & Hume, 1996) to obtain BP_ND_ values using PMOD’s PKIN module with a merged, bilateral cerebellum ROI serving as the reference region. Volume of interest (VOI)-based partial volume-corrected (PVC) estimates of BP_ND_ were computed based on work from Rousset, Ma, & Evans (1998) following the methods of Smith and colleagues (2017) which includes details on the specification of VOIs.

Mean BP_ND_ in the bilateral midbrain was extracted from an ROI drawn in MNI standard space using previously described guidelines (Dang et al., 2012, Dang et al., 2013) and registered to PET images using the same transformations for cerebellum registration to PET images. The midbrain ROI was not PVC. The PVC procedure requires specifying boundaries around a region and we did not collect an anatomical image that would have allowed the contrast required to precisely define these boundaries in the midbrain (e.g., GRASE or FFE; Eapen et al., 2011). For consistency with the earlier papers reporting evidence for the correlative triad that also used a whole striatum ROI, here, a whole striatum measure of PVC BP_ND_ was computed as a volume-weighted average of the caudate, putamen, and ventral striatum ROIs.

### Region-of-interest analyses

To limit the number of statistical tests, we created volume-weighted BP_ND_ summary measures for each ROI. We limited our analyses to regions with the highest averaged levels of uncorrected BP_ND_ exceeded group-averaged BP_ND_ that have been used in previous analyses of these datasets (e.g., Castrellon et al., 2019). For the following analyses, we used a whole striatum ROI consisting of caudate, putamen, and ventral striatum since much of the prior literature has investigated the correlative triad in the whole striatum. Data for striatal subregions is included on the OSF project page: https://osf.io/xjqe9/.

All statistical tests were conducted in R. Within each study, linear regression analyses were used to evaluate the correlative triad. First, regression analyses examined the effects of age on cognitive performance. Second, regression analyses examined the effects of D2R BP_ND_ on cognitive performance controlling for the linear effect of age. Separate regression analyses were run for each cognitive test and ROI. Both studies had ROI data for midbrain, anterior cingulate, thalamus, amygdala, hippocampus, and insula. Only Study 1 which used the tracer Fallypride included the striatum, because the FLB tracer used in Study 2 does not produce stable estimates of D2R BP_ND_ in striatum (Halldin et al., 1995). Given that previous studies reported significant associations between D2R BP_ND_ within many of these ROIs and some of these cognitive tests, we used a liberal statistical threshold of p < .05, two-tailed (Tables 1, 2) to indicate significant associations. We only conducted formal tests of mediation if two conditions were met: (1) age was negatively associated with cognitive performance and (2) D2R BP_ND_ was positively associated with cognitive performance after controlling for age. To assess mediation, we started by using an early approach that was standard in the field when the correlative triad was proposed: is the effect of age on cognition significant before but not after controlling for D2R BP_ND_? We also computed bootstrapped estimates of the indirect effect (5000 replications) to evaluate whether age-related decline in D2R BP_ND_ significantly carried the influence of age to cognitive performance.

**Table 2.** Region of interest analyses for Study 1. Correlations between cognitive test scores and D2R BP_ND_ controlling for age (N=83). BP_ND_ estimates are PVC except midbrain which is uncorrected. * p < .05 uncorrected

|  | Trails A | Trails B-A | Digit Span | Letter-Number<br>Sequencing | Verbal Paired<br>Associates Delayed<br>Recall |
| --- | --- | --- | --- | --- | --- |
| Midbrain | .197 [−0.0237,0.418] | .032 [−0.216,0.279] | .115 [−0.127,0.359] | −.104 [−0.314,0.106] | −.007 [−0.213,0.199] |
| Striatum | **.270\* [0.060,0.479]** | −.123 [−0.360,0.115] | **.241\* [0.010,0.471]** | .039 [−0.165,0.243] | .141 [−0.053,0.336] |
| Anterior<br>cingulate | .140 [−0.0564,0.337] | −.017 [−0.202,0.235] | −.007 [−0.223,0.208] | −.089 [−0.275,0.097] | .112 [−0.067,0.292] |
| Thalamus | .112 [−0.0876,0.312] | −.140 [−0.359,0.079] | .042 [−0.176,0.260] | −.063 [−0.251,0.126] | −.020 [−0.204,0.163] |
| Amygdala | .084 [−0.118,0.287] | .098 [−0.124,0.320] | .025 [−0.195,0.245] | −.149 [−0.337,0.039] | .039 [−0.146,0.224] |
| Hippocampus | .0294 [−0.169,0.228] | .028 [−0.191,0.246] | −.010 [−0.225,0.205] | −.090 [−0.275,0.095] | −.052 [−0.232,0.128] |
| Insula | **.239\* [0.038,0.441]** | .028 [−0.201,0.256] | .055 [−0.171, 0.285] | −.094 [−0.288, 0.101] | −.076 [−0.265,0.113] |

All data and code are publicly available on OSF: https://osf.io/xjqe9/

### Whole-Brain Analyses

To explore potential associations outside of the ROIs, whole-brain voxelwise analyses used uncorrected estimates of D2R BP_ND_ (that were not PVC). Whole-brain analyses controlled for multiple comparisons using threshold-free cluster enhancement with 5000 permutations per contrast computed using the “randomise” function in FSL. Positive and negative contrasts were computed for each of the cognitive tasks for each study while controlling for age. Significance was assessed using a 0.05 threshold on the corrected p maps. FLB BP_ND_ maps were smoothed at 8 mm prior to group-level analysis (Plaven-Sigray et al., 2017). If a region was identified that positively correlated with cognitive performance (i.e., higher BP_ND_ = better performance), we conducted formal tests of mediation in R using the same procedures as above.

All beta coefficients for linear effects are reported as standardized betas.

## Results

### Study 1

#### Effects of age on cognition

In Study 1, Trails A and Trails B-A were positively correlated with age, indicating longer time to completion as age increased (Trails A: β_81_ = 0.461, [0.268, 0.655], p = 0.0001, Trails B-A: β_81_ = 0.222, [0.009, 0.434], p = 0.044). The three memory-related tasks all negatively correlated with age (digit span total: β_81_ = –.274, [–0.483, – 0.064], p = .012, letter number sequencing: β_81_ = –.554, [–0.735, –0.372], p < 0.0001, delayed recall: β_77_ = –0.614, [–0.790, –0.438], p < 0.0001).

#### Effects of age on dopamine D2R availability

For five out of the seven ROIs in Study 1, D2R BP_ND_ was significantly negatively correlated with age (amygdala: β_81_ = –.292, [–0.500,– 0.084], p=0.007, thalamus: β_81_= –0.260, [–0.470,–0.049], p=0.018, midbrain: β_81_=–0.504, [– 0.693,–0.316], p<0.0001, striatum: β_81_= –.450, [–0.644,–0.255], p<0.0001, insula: β_81_= –0.359, [–0.562,–0.156], p<0.0001). D2R BP_ND_ in the hippocampal and the anterior cingulate ROIs did not significantly correlate with age (hippocampus: β_81_= –0.201, [–0.414, 0.012], p=0.068, anterior cingulate cortex: β_81_= –0.21, [–0.425, 0.001], p=0.054.

#### Effects of dopamine D2R availability on cognition

When controlling for age in Study 1, only striatal D2R BP_ND_ positively correlated with task performance such that higher binding was associated with better performance (for all correlations between D2R BP_ND_ and cognitive tasks see Table 2). Striatal D2R BP_ND_ was positively correlated with digit span total score (β_81_=0.24, 95% CI=[0.010, 0.471], p=0.044) suggesting that higher D2R BP_ND_ was associated with better task performance. Striatal D2R BP_ND_ and insula D2R BP_ND_ also positively correlated with length of time spent completing the Trails A task (striatum: β_81_=0.270, [0.060,0.479], p=0.014, insula: β_81_=0.239, [0.038, 0.440], p=0.023), a counterintuitive finding suggesting that higher D2R availability was associated with worse task performance. For associations between D2R availability and both cognition and age using uncorrected (non-PVC) BP_ND_, see Supplementary Tables 1–2. For associations between age and cognition controlling for D2R availability, see Supplementary Table 3.

The whole-brain analyses of the Fallypride data revealed several regions where higher D2R BP_ND_ was significantly associated with better memory on the verbal paired associates delayed recall test. There were positive correlations between delayed recall and D2R BP_ND_ along the medial frontal gyrus, within anterior portions of superior and middle frontal gyri, and in superior portions of the body of the caudate. We created spherical ROIs in each of the two frontal cortical regions to evaluate potential mediation. We used an existing caudate ROI that was estimated during the PVC procedure. Unthresholded statistical maps can be found on NeuroVault at: https://neurovault.org/collections/3707/.

#### Mediation Analyses

The correlation between striatal D2R BP_ND_ and digit span was the only effect within a priori ROIs to meet our criteria for mediation analyses. In Study 1, the effect of age on digit span was reduced from a significant effect of β = –.27 to a nonsignificant effect of β = –.17 after controlling for striatal D2R BP_ND_ (see Figure 1). Although 40% of the total effect was mediated by striatal BP_ND_, bootstrapped estimates of the indirect effect were not significant, a*b= – .024, 95% CI = [–.053, –.001], z = –1.861, p = .063, despite the modest effect size.

**Figure 1.**
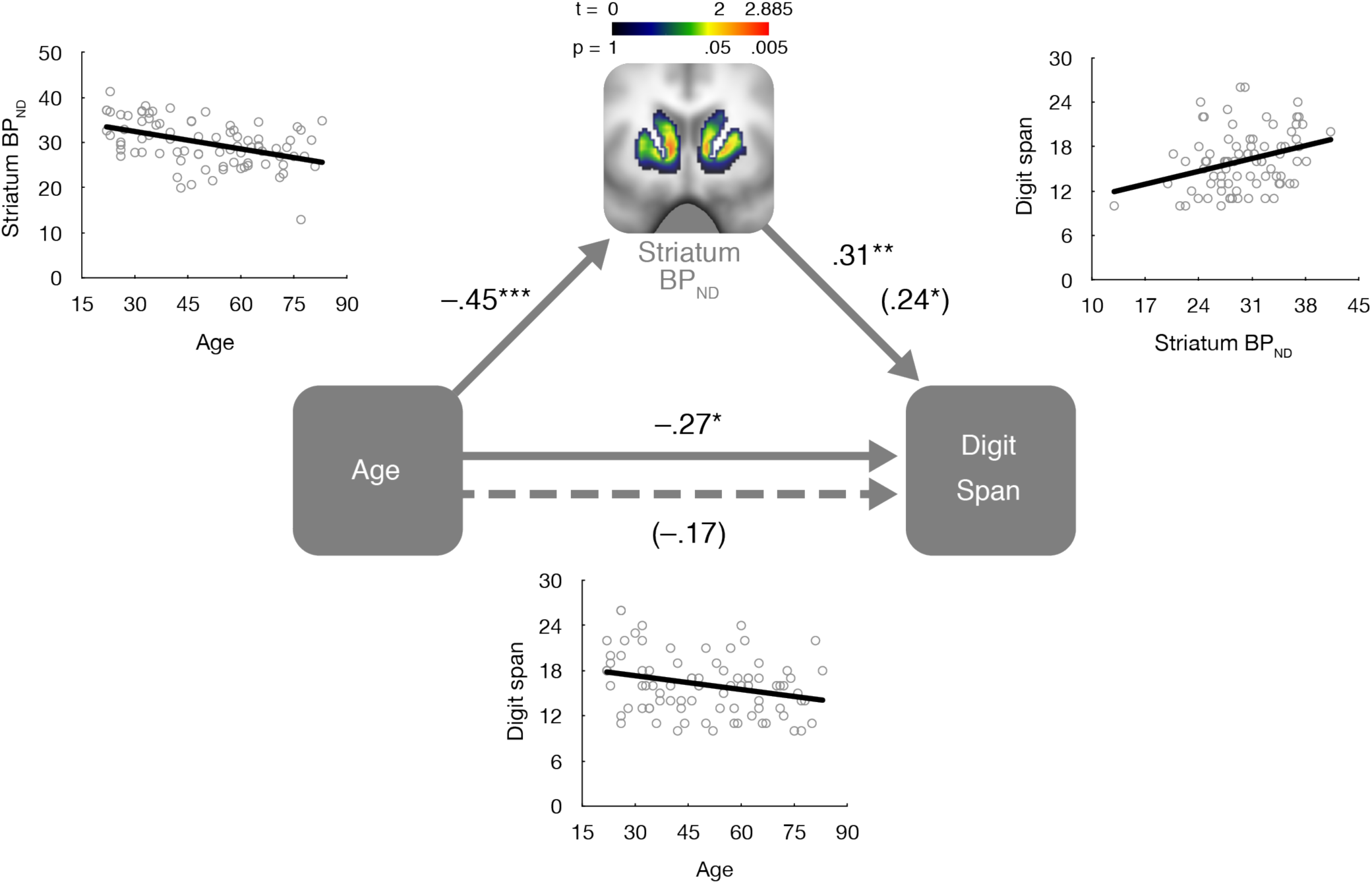
Associations between age, striatal dopamine D2R availability (BP_ND_), and digit span. Age significantly and negatively correlated with striatal dopamine D2R availability. Striatal dopamine D2R availability was significantly and positively correlated with digit span before controlling for age and after controlling for age (effect within parentheses). The brain image displays a voxelwise t-statistic map within the striatal ROI for visualization of regional variation in the association between individual differences in D2R availability and digit span. The effect of age on digit span was reduced from significant to nonsignificant (effect reported within parentheses) after controlling for striatal D2R availability. All scatterplots display simple pairwise associations between variables without covariates. Effect sizes displayed are standardized linear effects. * p < .05, ** p < .01, *** p < .001

We also evaluated mediation within three regions that were identified in whole brain analyses to be positively correlated with delayed recall for verbal paired associates (medial frontal gyrus, superior/middle frontal gyrus, superior caudate). Within all of these regions, the effect of age on delayed recall was significant before and after controlling for D2R BP_ND_ (medial frontal gyrus: β = –.61 to β = –.46; superior/middle frontal gyrus: β = –.61 to β = –.52; caudate: β = –.61 to β = –.57). Bootstrapped estimates of the indirect effect of D2R BP_ND_ were not significant in the superior/middle frontal gyrus, a*b=–0.012, 95% CI = [–0.033, 0.006], z=– 1.217, p = 0.224, medial frontal gyrus, a*b = –.02, 95% CI=[–0.054, 0.011], z=–1.244,p=0.214, or caudate, a*b= –.006, 95% CI=[–.022, .005],z= –.925, p=.355. Thus, none of the associations in these additional regions identified in the whole-brain analyses were consistent with statistical mediation by D2R availability of the effect of age on delayed recall.

### Study 2

#### Effects of age on cognition

In Study 2, Trails A and Trails B-A were significantly and positively correlated with age, indicating longer time to completion at older compared to younger ages (Trails A: β_35_=0.445, [0.149,0.742], p=0.006, Trails B-A: β_35_=0.499, [0.211,0.786], p=0.002). Long-term memory and one working memory task were significantly and negatively correlated with age (delayed recall: β_35_= –0.521, [–0.803, –0.238], p=0.001, letter number sequencing: β_35_= –0.393, [–0.697,–0.088], p=0.016). Digit span did not correlate with age (β_35_= –.070, [–0.401,0.260] p=0.678) in Study 2, although the confidence interval of the effect of age on digit span was highly overlapping with Study 1.

#### Effects of age on dopamine D2R availability

For four out of the six ROIs in Study 2, D2R BP_ND_ was significantly and negatively correlated with age (midbrain: β_35_= –0.640, [– 0.895,–0.386], p<0.0001, anterior cingulate cortex: β_35_= –0.477, [–0.769,–0.186], p=0.003, amygdala: β_35_= –0.340, [–0.652,–0.029], p =0.039, insula: β_35_= –0.438, [–0.736, –0.140], p=0.007). The hippocampal and thalamic ROIs did not have significant negative correlations between age and D2R BP_ND_ (thalamus: β_35_= –0.216, [–0.540,0.106], p=0.197, hippocampus: β_35_= –0.161, [–0.488,0.166], p=0.341).

#### Effects of dopamine D2R availability on cognition

When controlling for age in Study 2, letter number sequencing total score negatively correlated with D2R BP_ND_ in the midbrain (β_34_= –0.453, [–0.826,–0.081], p=0.023) and the hippocampus (β_34_=–0.323, [–0.617,–0.030], p=0.038), (for all correlations between task and D2R BP_ND_ see Table 3) such that higher D2R BP_ND_ was associated with worse performance. Likewise, Trails B-A was positively correlated with D2R BP_ND_ in the midbrain (β_34_= 0.390, [0.034,0.746], *p*=0.039) such that higher D2R availability was associated with worse performance. For associations between D2R availability and both cognition and age using uncorrected (non-PVC) BP_ND_, see Supplementary Tables 4–5. For associations between age and cognition controlling for D2R availability, see Supplementary Table 6.

**Table 3.** Region of interest analyses for Study 2. Correlations between cognitive test scores and D2R BP_ND_ controlling for age (N=37). BP_ND_ estimates are PVC except midbrain which is uncorrected. * p < .05 uncorrected

|  | Trails A | Trails B-A | Digit Span | Letter-Number<br>Sequencing | Verbal Paired<br>Associates Delayed<br>Recall |
| --- | --- | --- | --- | --- | --- |
| Midbrain | -.289 [-0.669,0.090] | **.390\* [0.0341, 0.746]** | -.376 [-0.794,0.0416] | **-.453\* [-0.826,-0.081]** | -.169 [-0.538,0.200] |
| Anterior<br>cingulate | -.008 [-0.350,0.335] | .189 [-0.136,0.515] | -.076 [-0.457,0.305] | -.184 [-0.530,0.163] | -.048 [-0.374,0.278] |
| Thalamus | -.092 [-0.399,0.215] | .213 [-0.076,0.503] | -.146 [-0.486,0.194] | -.226 [-0.534,0.081] | .136 [-0.155,0.426] |
| Amygdala | -.191 [-0.505,0.122] | .199 [-0.103,0.502] | -.080 [-0.436,0.275] | -.313 [-0.624,-0.001] | -.155 [-0.456,0.146] |
| Hippocampus | .012 [-0.292,0.317] | .140 [-0.151,0.431] | -.143 [-0.479,0.193] | **-.323\* [-0.617,-0.030]** | .082 [-0.208,0.371] |
| Insula | .016 [-0.312,0.351] | .309 [0.002, 0.616] | -.183 [-0.551,0.184] | -.187 [-.525,0.151] | -.081 [-0.399,0.237] |

The whole-brain analyses of the FLB data did not reveal any regions where higher D2R BP_ND_ was significantly associated with better performance. There were negative correlations with letter-number sequencing and D2R BP_ND_ in lateral temporal cortex and lateral occipital cortex. There were positive correlations with Trails B-A and D2R BP_ND_ in superior and middle temporal gyrus. These effects were opposite of the predicted direction such that higher D2R availability corresponded to worse performance. Thus, we did not evaluate mediation for any of these effects within the FLB data. Unthresholded statistical maps can be found on NeuroVault at: https://neurovault.org/collections/3707/.

Given the positive associations between delayed recall and Fallypride D2R BP_ND_ in Study 1, we also evaluated associations between memory tasks and FLB D2R BP_ND_ and potential mediation by FLB D2R BP_ND_ in Study 2 within the two regions identified in Study 1 that were available for analysis in Study 2 (medial frontal gyrus, superior/middle frontal gyrus). D2R BP_ND_ was not correlated with delayed recall in either of these regions (medial frontal gyrus: β=– .11; superior/middle frontal gyrus: β=.07). Given the lack of association between D2R BP_ND_ and delayed recall there was an expected lack of a mediation by D2R BP_ND_. The effect of age on delayed recall was significant before and after controlling for D2R BP_ND_ (medial frontal gyrus: β=–.52^z^ to β=–.60; superior/middle frontal gyrus: β=–.52 to β=–.48); bootstrapped estimates of the indirect effect were not significant in the medial frontal gyrus, a*b=–0.013, 95% CI=[–.024, .055], z=.658, p=.51, or superior/middle frontal gyrus, a*b=–0.007, 95% CI=[–.055, .022], z=– .388, p=.698. Overall the associations between cortical D2R availability and memory were not consistent across studies using different tracers (Study 1: Fallypride; Study 2: FLB) and neither set of results provided evidence for mediation.

## Discussion

Here, in two cross-sectional studies of adulthood, we examined associations between aging, dopamine D2R availability using PET imaging, and cognition which form the bases for the correlative triad theory. We found that very few associations were significant and only one was in the predicted direction. While associations between aging and cognition and aging and D2R availability were mostly significant and highly reproducible across studies, the only region and measure that was consistent with the correlative triad was striatal D2R BP_ND_ and digit span in Study 1 that used Fallypride. Although inclusion of the dopamine D2R measure reduced the age effect on digit span from significant to non-significant, statistical evaluation of mediation was non-significant. Our findings provide partial support for the correlative triad in one data set for working memory, but qualify general claims of an association between age-related decline in the dopamine system and age-related decline in fluid cognition.

Several individual prior studies have reported associations between D2R availability across a variety of different cognitive measures. As a result, we did not have strong hypotheses about which of the cognitive measures would show the strongest effects. Consistent with the positive associations reported here between D2R availability and short-term maintenance, several prior studies reported associations between dopamine target measures and short-term maintenance and/or working memory (positive: Erixon-Lindroth, et al., 2005 (dopamine transporter); Backman, et al., 2011 (D1); Salami, et al., 2018 (D2R); negative: Reeves, 2005 (D2R)). Two of the cognitive measures collected in the present studies assessed aspects of working memory: digit span total score and letter-number sequencing. Digit span, but not letter-number sequencing, was negatively associated with age and positively associated with striatal D2R availability in Study 1 using Fallypride. It is unclear why there was an association with digit span but not letter-number sequencing, given prior evidence that both tasks load onto the same latent factor (Parmenter, Shucard, Benedict, & Shucard, 2006), although it may be noted that letter number sequencing requires a more complex manipulation of information in working memory. One previous study indicated that digit span and logical memory were most strongly correlated with D2R availability when controlling for age (Chen et al., 2005). However, the majority of prior studies reporting associations between measures of dopamine targets (receptors, transporters, or relevant enzymes) and working memory used n-back tasks (see Salami et al., 2018). Interestingly, some research suggests that digit span does not strongly correlate with n-back, although these studies did not specifically use the combined forward and backward score used in the present studies (Miller, Price, Okkun, Montijo, & Bowers, 2009; Parmenter, Shucard, Benedict, & Shucard, 2006). Nevertheless, it is possible that we would have detected stronger associations between D2R availability and working memory if we used an n-back task. That said, Nyberg et al. (2016) did not find a relationship between working memory (and perceptual speed) and D2R availability measured using the tracer [^11^C]Raclopride, prompting a hypothesis that D2R might be more strongly related with episodic memory and the hippocampus.

We observed positive associations between digit span and D2R availability in the striatum but not in any other regions of interest. Although task-based neuroimaging studies most often highlight associations between prefrontal cortical function and working memory (Braver et al., 1997; Wager & Smith, 2003; Constantinidis & Klingberg, 2016), striatal dopamine synthesis capacity has also been associated with working memory (Landau, 2009). Additionally, lesions to the striatum produce deficits in working memory and many other fluid cognitive tasks assumed to be primarily dependent on the prefrontal cortex (Rubin, 1999). Furthermore, a pharmacological fMRI study showed that amphetamine administration increased striatal as well as frontal cortical BOLD signal during an n-back working memory task in older and younger adults (Garrett et al. 2015). The lack of associations between frontal D2R availability reported here might reflect the lower signal in lateral cortical regions for the radiotracers used in this study, particularly for Fallypride (Mukherjee et al., 2002; Vandehey et al., 2010; Zald et al., 2010). Estimates of frontal D2R availability using FLB are higher compared to Fallypride, but many lateral regions still have quite low signal, which may constrain the variance available to explain variation in cognition. Thus, it is possible that there are positive associations between D2R availability and cognition in regions of the frontal cortex that we were not able to detect. While we did observe an association between striatal D2R availability and digit span, this could only be tested within Study 1 using Fallypride (since FLB used in Study 2 does not produce stable estimates of striatal BP_ND_). Thus, we could not evaluate the replicability of that effect across data sets. Differences in tracer pharmacokinetics might also contribute to differences in the size of observed associations. The present studies utilized different tracers (i.e., the high affinity tracers Fallypride and FLB457) compared to the foundational studies in this area (which mostly used Raclopride), but we would not expect this to lead to a fundamentally different pattern of associations.

Exploratory whole-brain analyses identified additional frontal cortical regions that were positively associated with delayed recall in Study 1 using Fallypride. However, additional analyses provided no evidence for statistical mediation. In fact, the effects of age on delayed recall were nearly the same before or after controlling for D2R availability in these frontal regions in both studies. Finally, the associations between frontal D2R availability and delayed recall were not significant in Study 2 that used FLB which produces higher estimates of cortical D2R availability. Overall, there was no evidence for frontal cortical mediation within Study 1 using Fallypride or Study 2 using FLB and the associations between dopamine D2R availability and delayed recall were inconsistent across Studies 1 and 2.

Contrary to our hypotheses and the correlative triad theory, we also observed some *negative* associations between dopamine D2R availability and cognition such that higher D2R availability was associated with worse task performance. Other studies have also reported such negative associations. For example, in a recent study of a cohort of older adults, one subgroup had lower D2R availability in the striatum but also had higher working memory scores (Lövdén et al. 2017). Levels of D2R availability could reflect compensatory changes in response to lower endogenous DA levels. Thus, it is possible that the observed negative associations are indicators that relatively higher endogenous dopamine levels are associated with better cognitive performance. However, the consistent and highly replicable evidence for lower D2R availability in older age and lower fluid cognitive performance in older age runs counter to this interpretation especially in cross-sectional life-span studies. Note, also, that these effects were not consistent across samples within regions that were reported across both studies.

Though studies that laid the foundation for the correlative triad theory also used D2R tracers, it is possible that other components of dopamine function, such as other receptor subtypes or dopamine release, are better correlated with fluid cognitive ability. For example, D1-like receptor availability has been associated with motor performance (Wang et al., 1998) as well as several memory measures (Sawaguchi & Goldman-Rakic, 1991; Sawaguchi, 2001; Karlsson, 2009, Backman, 2011). However, even studies with other measures of dopamine function have failed to replicate some of these associations. For example, Rieckmann et al., 2018 found no significant associations between dopamine transporter density in the striatum and processing speed or executive function (factors composing some of the tests used in the present study). However, as noted earlier it is difficult to assess the reproducibility of these effects given the limited number of studies and inconsistent use of tasks.

Overall, many of the individual previously reported associations between dopamine D2R availability and cognition in smaller samples were not replicated in the present studies (e.g., Bäckman, 2000; Erixon-Lindroth et al., 2005). It is possible that the neuropsychological test measures used in the present studies lacked sensitivity relative to the tasks used in prior studies. Some prior studies reported associations between measures of dopamine function and measures of memory or cognitive flexibility based on scores from experimental behavioral tasks rather than neuropsychological measures. It is possible that those tasks better assess aspects of cognitive function that are most related to dopamine function. One possibility is that the neuropsychological tests in these two studies were not difficult enough. Salami and colleagues (2019) found that D2R-BOLD associations were greatest in a 3-back task compared to 1 or 2-back in older adults who performed normally on the working memory task, suggesting a relationship between D2R availability and task efficiency. Additionally, recent work suggests that individual differences in gene expression moderates the relationship between D2R availability and cognitive test scores (Karalija et al., 2019). Thus, associations between D2R availability and cognitive measures in the present studies may be masked by genetic variation, which has contributed to a more recent refinement of the correlative triad hypothesis (Karalija et al., 2019). Relatedly, other individual difference factors (e.g., education and body-mass index) modify associations between dopamine and cognition in initial analyses of the COBRA longitudinal dataset (Lovden et al., 2017). Given that the emerging moderating effects are based on statistical interactions and multivariate associations which require much more statistical power than simple main effects, future studies should evaluate the replicability of these more complicated effects. While both of the studies reported here, particularly Study 1, were larger than the majority of studies that originally explored the correlative triad, they were underpowered to investigate individual differences independent of age or potential interactions between age or D2R availability and other variables of interest. If others would like to conduct exploratory analyses of potential interactions with some of these variables, we also include sex, education, income, race/ethnicity, and BMI in our publicly available data files (https://osf.io/xjqe9/). While we have not ourselves tested for such moderating effects, the present studies could be consistent with the recent evidence for moderating factors given the lack of support for a broad traditional linear mediation model of the correlative triad.

Recent research has identified broad variation in the effects of age on D2R availability across subregions of the brain (Seaman et al., 2019). It is possible that there are subregional effects consistent with the correlative triad that we missed in the present analyses. The studies here focused on a limited set of somewhat larger ROIs. Many of these ROIs were the same size (e.g., whole striatum) or even smaller than what has been used in previous studies of the correlative triad. The primary goal of the present analyses was to evaluate reproducibility so we used ROIs that would be most consistent with previously published research. The whole-brain analyses were included to identify potential subregional effects, although no clear subregional effects emerged from these analyses.

A final and critical limitation of our findings, and the literature in general, is that despite our relatively large samples, conclusions about aging are drawn from cross-sectional data. It is challenging to accurately evaluate statistical mediation of aging from cross-sectional data. Mediation in cross-sectional data may not generalize in longitudinal samples, and a mediation effect in a longitudinal sample may exist despite a lack of significant mediation in cross-sectional data (Raz & Lindenberger, 2011). As in any cross-sectional study, the results reported here may be related to cohort rather than aging effects. Ongoing longitudinal work will be able to better address potential longitudinal associations between the dopamine system and cognition (Nevalainen et al., 2015).

Overall, the present findings suggest that associations between age differences in dopamine D2R availability and fluid cognition may not be as replicable or broad as initially assumed. There was some suggestive evidence for an association between D2R availability and short-term memory maintenance, but the majority of previously reported findings were not replicated here. A wealth of other research in human and non-human animals demonstrates that dopamine function makes critical contributions to cognition, but the present studies suggest caution in using PET data to make generalizations that dopamine receptor loss is a critical cause of broad age-related declines in fluid cognition.

## Acknowledgements

This research was supported by National Institute on Aging Pathway to Independence Award R00-AG042596, National Institute on Aging grant R01-AG044838, and National Center for Advancing Translational Science grant UL1TR000445. K.L.S. was supported by T32-AG000029.

## Data

Data used in the manuscript can be viewed and downloaded from: https://osf.io/xjqe9/

## SUPPLEMENTARY MATERIALS

**Supplementary Figure 1.**
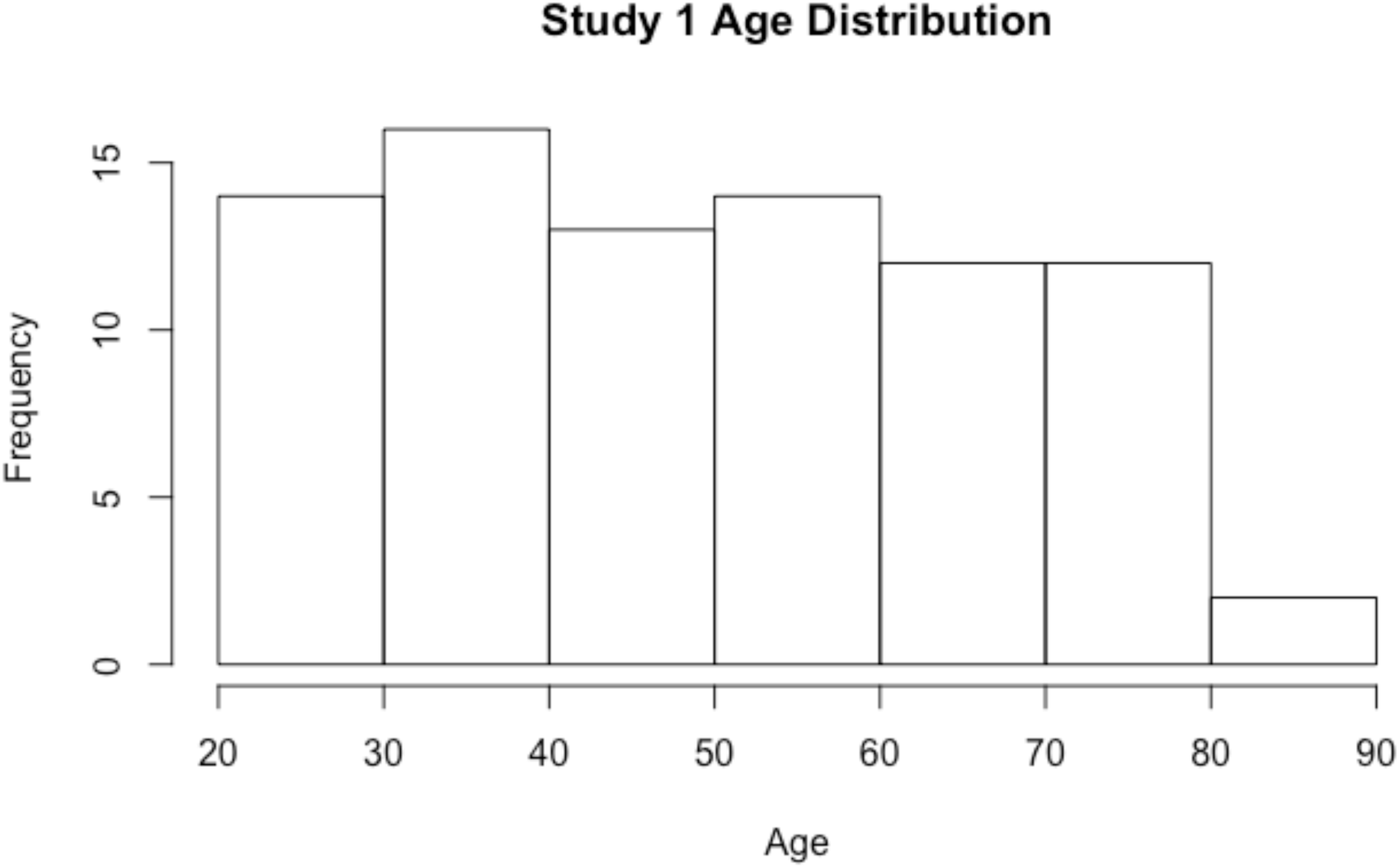
Age distribution for Study 1, N=83.

**Supplementary Figure 2.**
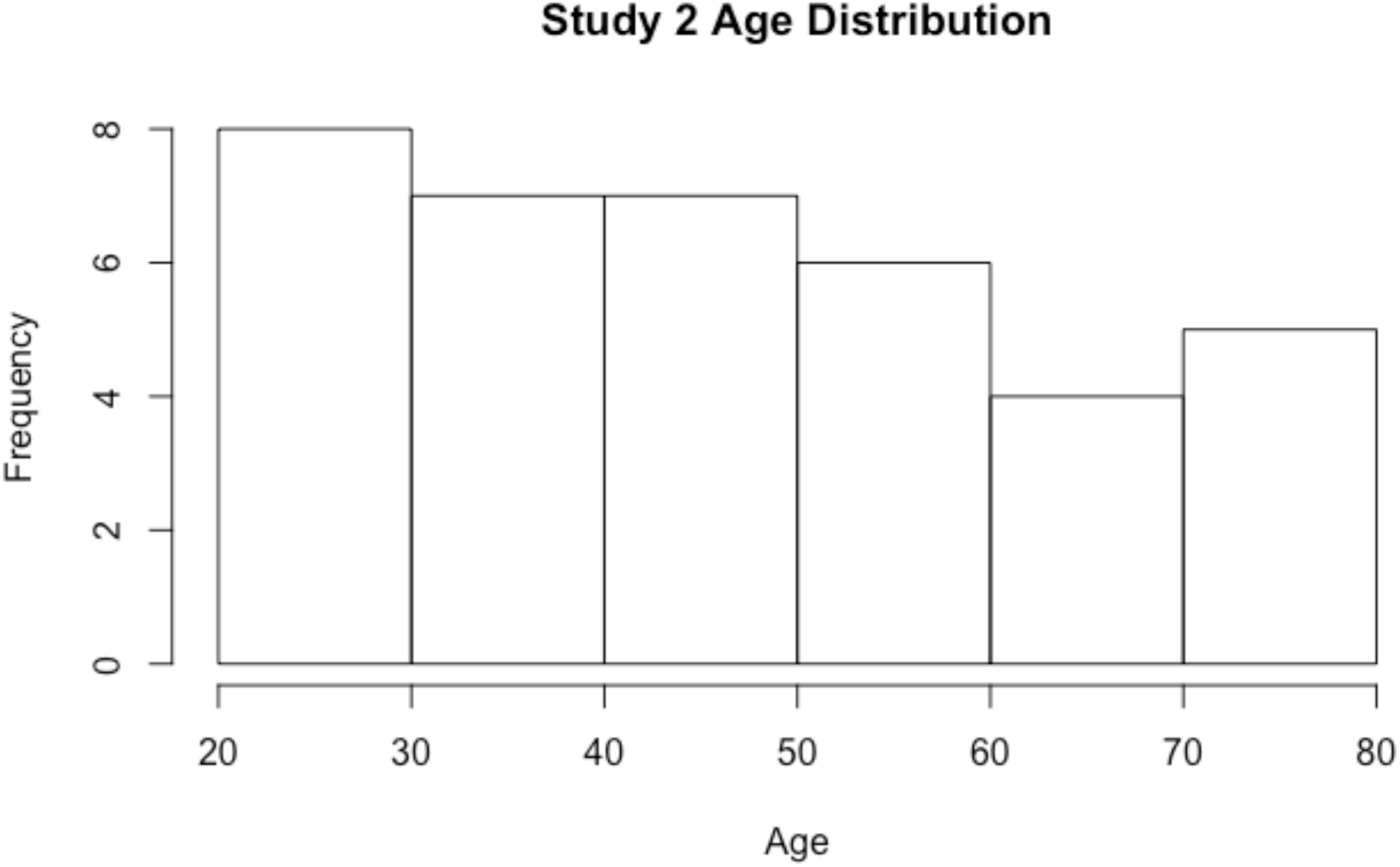
Age distribution for Study 2, N=37.

**Supplementary Table 1.** Non-PVC region of interest analyses for Study 1. (Note: midbrain ROI reported within the paper is already non-PVC.) Correlations between cognitive test scores and D2R BP_ND_ controlling for age (N=83). * p < .05 uncorrected

|  | Trails A | Trails B-A | Digit Span | Letter-Number Sequencing | Verbal Paired Associates Delayed Recall |
| --- | --- | --- | --- | --- | --- |
| Striatum | **.277\* [0.059,0.495]** | -.131 [-0.378,0.116] | **.240\* [0.001,0.480]** | .033 [-0.179,0.246] | .141 [-0.061,0.343] |
| Anterior cingulate | **.221\* [0.014,0.428]** | -.154 [-0.385,0.078] | .007 [-0.224,0.238] | -.124 [-0.322,0.073] | .002 [-0.191,0.195] |
| Thalamus | .105 [-0.100,0.310] | -.132 [-0.358,0.093] | .044 [-0.179,0.268] | -.063 [-0.256,0.131] | -.016 [-0.204,0.173] |
| Amygdala | .087 [-0.122,0.296] | .101 [-0.129,0.330] | .006 [-0.234,0.222] | -.107 [-0.302,0.089] | .008 [-0.183,0.199] |
| Hippocampus | .064 [-0.136,0.264] | .006 [-0.215,0.226] | -.003 [-0.243,0.191] | -.114 [-0.301,0.072] | -.072 [-0.254,0.109] |
| Insula | **.239\* [0.023,0.454]** | .018 [-0.226,0.262] | .091 [-0.149, 0.331] | -.065 [-0.273, 0.143] | -.062 [-0.264,0.139] |

**Supplementary Table 2.** Non-PVC region of interest analyses for Study 1. (Note: midbrain ROI reported within the paper is already non-PVC.) Correlations between age and D2R BP_ND_ (N=83). * p < .05 uncorrected

|  | Age |
| --- | --- |
| Striatum | **-0.511\*, [-0.698, -0.323], p < .0001** |
| Anterior cingulate | **-0.407\*, [-0.606, -0.208], p = .0001** |
| Thalamus | **-0.339\*, [-0.544, -0.134], p = .002** |
| Amygdala | **-0.378\*, [-0.580, -0.177], p = .0004** |
| Hippocampus | **-0.247\*, [-0.458, -0.036], p = .024** |
| Insula | **-0.482\*, [-0.673, -0.292], p < .0001** |

**Supplementary Table 3.** Correlations between age and cognitive tests before (first row after *Simple effect of age*) and after controlling for D2R BP_ND_ in Study 1 (N=83) *p<.05 uncorrected

|  | Trails A | Trails B-A | Digit Span | Letter-Number Sequencing | Verbal Paired Associates Delayed Recall |
| --- | --- | --- | --- | --- | --- |
| Simple effect of age | **.461\* [0.268, 0.665]** | **.222\* [0.009, 0.434]** | **-.274\* [-0.483, -0.064]** | **-.554\* [-0.735, -0.372]** | **-.614\* [-0.790, -0.438]** |
| Age effect controlling for regional D2 BP<sub>ND</sub> in... |  |  |  |  |  |
| Midbrain | **.561\* [0.340, 0.782]** | .238 [-0.009, 0.485] | -.215 [-0.458, 0.028] | **-.606\* [-0.816, -0.396]** | **-.618\* [-0.824, -0.411]** |
| Striatum | **.583\* [0.373, 0.792]** | .167 [-0.071, 0.404] | -.165 [-0.395, 0.065] | **-.536\* [-0.740, -0.332]** | **-.552\* [-0.747, -0.358]** |
| Anterior cingulate | **.491\* [0.294, 0.688]** | **0.225\* [0.007, 0.444]** | **-.275\* [-0.491, -0.059]** | **-.572\* [-0.758, -0.387]** | **-.590\* [-0.770, -0.411]** |
| Thalamus | **.490\* [0.291, 0.690]** | .186 [-0.034, 0.405] | **-.263\* [-0.481, -0.045]** | **-.570\* [-0.758, -0.381]** | **-.619\* [-0.803, -0.436]** |
| Amygdala | **.486\* [0.284, 0.688]** | **.250\* [0.028, 0.473]** | **-.027\* [-0.487, -0.046]** | **-.597\* [-0.785, -0.409]** | **-.603\* [-0.788, -0.418]** |
| Hippocampus | **.467\* [0.269, 0.666]** | **.227\* [0.009, 0.445]** | **-.276\* [-0.491, -0.060]** | **-.572\* [-0.757, -0.387]** | **-.624\* [-0.804, -0.443]** |
| Insula | **.547\* [0.346, 0.749]** | .232 [0.003, 0.461] | **-.254\* [-0.479, -0.029]** | **-.587\* [-0.782, -0.393]** | **-.641\* [-0.830, -0.452]** |

**Supplementary Table 4.**
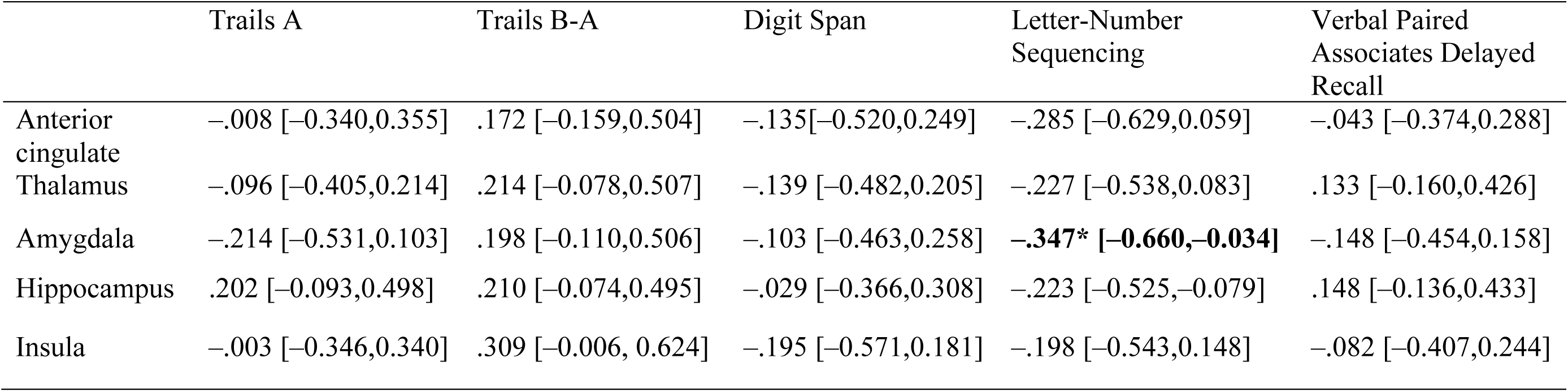
Non-PVC region of interest analyses for Study 2. (Note: midbrain ROI reported within the paper is already non-PVC.) Correlations between cognitive test scores and D2R BP_ND_ controlling for age (N=37). * p < .05 uncorrected

**Supplementary Table 5.** Non-PVC region of interest analyses for Study 2. (Note: midbrain ROI reported within the paper is already non-PVC.) Correlations between age and D2R BP_ND_ (N=37). * p < .05 uncorrected

|  | Age |
| --- | --- |
| Anterior cingulate | **-0.500\*, [-0.787, -0.213], p = .002** |
| Thalamus | -0.253, [-0.573, 0.068], p = .132 |
| Amygdala | **-0.378\*, [-0.685, -0.072], p = .021** |
| Hippocampus | -0.112, [-0.441, 0.218], p = .511 |
| Insula | **-0.479\*, [-0.770, -0.188], p = .003** |

**Supplementary Table 6.** Correlations between age and cognitive tests before (first row after *Simple effect of age*) and after controlling for D2R BP_ND_ in Study 2 (N=37) *p<.05 uncorrected

|  | Trails A | Trails B-A | Digit Span | Letter-Number Sequencing | Verbal Paired Associates Delayed Recall |
| --- | --- | --- | --- | --- | --- |
| Simple effect of age | **.445\* [0.149, 0.742]** | **.499\* [0.211,0.786]** | -.070 [-0.401,0.260] | **-.393\* [-0.697,-0.088]** | **-.521\* [-0.803,-0.238]** |
| Age effect controlling for regional D2 BP<sub>ND</sub> in... |  |  |  |  |  |
| Midbrain | .260 [-0.120,0.640] | **.748\* [0.392,1.104]** | -.311 [-0.729,0.106] | **-.683\* [-1.055,-0.310]** | **-.629\* [-0.998,-0.260]** |
| Anterior cingulate | **.442\* [0.099,0.784]** | **0.589\* [0.263, 0.914]** | -.107 [-0.488,0.274] | **-.480\* [-0.827,-0.134]** | **-.544\* [-0.870,-0.217]** |
| Thalamus | **.425\* [0.118,0.732]** | **.545\* [0.255,0.835]** | -.102 [-0.442,0.238] | **-.442\* [-0.749,-0.134]** | **-.491\* [-0.782,-0.201]** |
| Amygdala | **.380\* [0.067,0.694]** | **.566\* [0.264,0.869]** | -.098 [-0.453,0.258] | **-.499\* [-0.810,-0.188]** | **-.573\* [-0.874,-0.273]** |
| Hippocampus | **.447\* [0.142,0.752]** | **.521\* [0.230,0.813]** | -.094 [-0.430,0.243] | **-.445\* [-0.738,-0.151]** | **-.507\* [-0.800,-0.218]** |
| Insula | **.452\* [0.117,0.787]** | **.634\* [0.327,0.941]** | -.151 [-0.519, 0.217] | **-.474\* [-0.812, -0.136]** | **-.556\* [-0.874,-0.238]** |

